# *NAT8L* mRNA oxidation is linked to neurodegeneration in multiple sclerosis

**DOI:** 10.1101/2020.04.19.049494

**Authors:** Prakash Kharel, Naveen Kumar Singhal, Nicole West, Joram Rana, Lindsey Smith, Ernest Freeman, Ansuman Chattopadhyay, Jennifer McDonough, Soumitra Basu

## Abstract

RNA oxidation has been implicated in neurodegeneration, but the underlying mechanism for such effects is unclear. Recently, we demonstrated extensive RNA oxidation within the neurons in multiple sclerosis (MS) brain. In this report we identified selectively oxidized mRNAs in neuronal cells that pertained to neuropathological pathways. N-acetyl aspartate transferase 8 like (*NAT8L*) mRNA is one such transcript, whose translated product enzymatically synthesizes N-acetyl aspartic acid (NAA), a neuronal metabolite important for myelin synthesis. We reasoned that impediment of translation of an oxidized *NAT8L* mRNA will result in reduction in its cognate protein, thus lowering NAA level. This assertion is directly supported by our studies on a model cellular system, an MS animal model and postmortem human MS brain. Reduced NAA level in the brain hampers myelin integrity making neuronal axons more susceptible to damage, which contributes in MS neurodegeneration. Overall, this work provides a framework for mechanistic understanding of the link between RNA oxidation and neurodegenerative diseases.

## Introduction

Multiple sclerosis (MS) is the most common inflammatory and demyelinating disorder of the central nervous system (CNS), affecting more than 2.3 million people globally (1). The degeneration and loss of neuronal axons are the major causes of the progressive neurological decline, a symptom found in the majority of MS patients. In addition to inflammation, oxidative stress and mitochondrial dysfunction contribute to MS neurodegeneration (2-4). The inadequacy of free radical scavenging machinery in the cells results in the overproduction of reactive oxygen species (ROS), which can oxidatively modify biomolecules including proteins, lipids, and nucleic acids (5). A higher level of lipid peroxidation, protein nitration, and nucleic acids oxidation has been reported in the different areas of postmortem MS brain (2, 6-9). Although, a significant effort has been invested to understand the impact of major biomolecules on the process of neurodegeneration, deciphering the effect of RNA oxidation has been largely overlooked. Recently, RNA oxidation has been proposed as a contributing factor to the neurodegeneration in many neurodegenerative diseases including Alzheimer’s disease (AD) (10-12), amyotrophic lateral sclerosis (ALS) (13), Parkinson’s disease (PD) (14), and MS (6). While the link between RNA oxidation and neurodegeneration in several neurodegenerative diseases has been widely speculated (6, 10, 13, 15, 16), detailed studies about the identity of the oxidized RNAs and their role in the molecular mechanisms resulting in pathogenesis are limited.

In RNA, guanosine is the most oxidizable nucleoside that produces the mutagenic product 8-oxoguanosine (8-OG, also widely referred to as 8-hydroxyguanosine, 8-OHG based on the less stable tautomer) upon oxidation (17, 18). Besides base pairing with its canonical partner cytosine, the oxidation product 8-OG can non-canonically base pair with adenine to cause mutagenic effects (19). The presence of 8-OG in the open reading frame (ORF) of mRNA has shown to result in truncated proteins (20) and ribosome stalling (21). The studies also showed that oxidized mRNAs have a similar lifetime as the non-oxidized mRNAs *in cellulo*, which may indicate the lack of a global RNA 8-OG editing mechanism, in turn suggesting that oxidized mRNAs can contribute to cellular pathophysiology. Oxidatively damaged bases are repaired in the DNA and there are speculations about the editing of such RNA lesions (22). A recent report showed that binding of the PCBP1 protein to oxidized RNA induces cell death (23). However, the existence of global RNA oxidative modification editing mechanism has not yet been identified. Oxidized transcripts can have role in the pathogenesis of neurological disorders, such as, MS.

Deficiency of N-acetyl aspartate (NAA) is associated with MS neurodegeneration. N-Acetyl aspartate transferase 8 like protein (NAT8L) is a key trans-mitochondrial membrane protein that catalyzes the synthesis of NAA, which is one of the most abundant analytes (∼20 mM in neurons) in the central nervous system (CNS) (24-26). Once synthesized in the neurons, NAA is either exported to the oligodendrocytes or used in N-acetylaspartylglutamate (NAAG) synthesis within the neurons to be exported to astrocytes (27). Once aspartoacylase (ASPA) hydrolyzes NAA to aspartate and acetyl-CoA in the oligodendrocytes, the acetate moiety from acetyl-CoA becomes involved in myelin synthesis, which is deposited on neuronal axons as a protective insulating material (28). Interestingly, it has been reported that NAA-derived acetate in the oligodendrocytes has a three-fold higher potential to be incorporated into the myelin lipids than the free acetate (29). In addition to NAA’s role in myelin synthesis in oligodendrocytes, NAA has been proposed to act as an organic osmolyte that removes excess water from neurons by acting as a molecular water pump (30). The proposed roles of NAA also include the maintenance of intact white matter, promotion of dopamine uptake by regulating TNF-alpha expression, and attenuation of methamphetamine-induced inhibition of dopamine uptake (31). The McDonough lab discovered that NAA regulates histone H3 methylation in oligodendrocytes and also lipid composition and structure of myelin (32). Sumi et al. observed that axonal degeneration in the MS brain was correlated with a reduced level of NAA (33). Reduced NAA levels could reflect reversible neuronal damage caused by inflammatory demyelination, altered neuronal metabolism, or axonal loss (33). *Nat8l* knockout mice (*Nat8l*^−/−^) showed decreased NAA content in the brain, less compact myelin structure, reduced social interaction, and shortened immobility time in the forced swimming test (34). As demonstrated by magnetic resonance spectroscopy (MRS) studies on patients (33) and HPLC analysis of NAA in the postmortem MS brain extracts (32, 35), the reduction of NAA is one of the pathological hallmarks in the CNS of MS patients. Although, the reduction of NAA in the MS brain has detrimental effects (32), the causative factor behind low levels of NAA is still unclear.

We reasoned that oxidative stress induced selective mRNA oxidation in neurons could be linked to MS pathogenesis. Using RNA-seq and RT-qPCR analyses of the oxidized mRNA pool from human cultured neurons, we identified the selectively oxidized mRNAs under conditions that reflect the MS microenvironment. We discovered that *NAT8L* mRNA was one of the selectively oxidized mRNAs that led us to hypothesize that oxidized *NAT8L* mRNA could result in reduced NAT8L protein production causing lowered NAA levels, which may contribute to MS pathogenesis. Indeed, the oxidized *NAT8L* mRNA resulted in the reduced production of NAT8L protein *in vitro*. We also confirmed these changes in the cuprizone mouse model of MS (36).

Cuprizone mice mimic many aspects of MS pathology. Cuprizone feeding induces oligodendrocyte apoptosis, demyelination, and activation of microglia that generate increased reactive nitrogen species (RNS). The fact that these mice also have reduced levels of NAA make them appropriate for this study (37, 38). Taken together, the *in cellulo* and *in vivo* studies under oxidative stress revealed that a higher *NAT8L* mRNA oxidation leads to a reduced production of NAT8L protein in the MS microenvironment causing lowered NAA levels. Our study established a molecular link between selective mRNA oxidation and MS pathogenesis.

## Results

### *NAT8L* mRNA is selectively oxidized in human neurons under oxidative stress

Recently, we discovered that mRNAs are highly oxidized in the neurons within the normal appearing areas of MS brain (6). Activated microglia mediated production of nitroxide (NO^**·**^) and hydroxide (^**·**^OH) radicals are considered key to MS pathogenesis (39). Using sodium nitroprusside (SNP), an NO^**·**^ and ^**·**^OH donor molecule (Figure 1a) (40), we induced a mild oxidative stress in SH-SY5Y human neuroblastoma cells. The oxidative stress (OS) in the cells treated with SNP was verified by quantification of the H_2_O_2_ produced (Figure 1b). Next, the relative impact of OS on mRNA was quantified by comparing the relative amount of 8-OG present in the total cellular RNA with or without SNP treatment. The SNP treated cells showed a three-fold increase in oxidized mRNA compared to the non-treated cells. To decipher the identity of oxidatively modified mRNAs, the oxidized mRNA pool was isolated by immunoprecipitation with anti 8-OG antibody followed by oligo-dT mRNA enrichment from the SNP treated SH-SY5Y cells (schematic illustration in Figure 1c). After the selection of the oxidized mRNA pool, we ensured that there was a greater extent of oxidative damage in this immunoprecipitated mRNA pool using HPLC-ECD detection of 8-OG, the major oxidation product. We found a three-fold increase in mRNA oxidation in the SNP treated cells (SI Appendix, Figure S1). The quality of the immunoprecipitated mRNA was tested by Bioanalyzer mRNA integrity test before sequencing (Girihlet Inc.) (Figure 1c). iLumina^®^ RNA-seq platform was used to investigate the identity of the oxidized mRNAs and the differential upregulation of oxidized mRNA in the SNP treated neuronal cells was calculated in comparison to the control mRNA pool. The goal of RNA-seq analysis of oxidized mRNA in SNP treated cells was to determine whether MS related transcripts were selectively oxidized in the neuronal cells. Indeed, we observed 819 transcripts were more abundant (fold change ≥ 1.5, P ≤ 0.05) in the oxidized mRNA pool in comparison to the control RNA.

**Figure 1.**
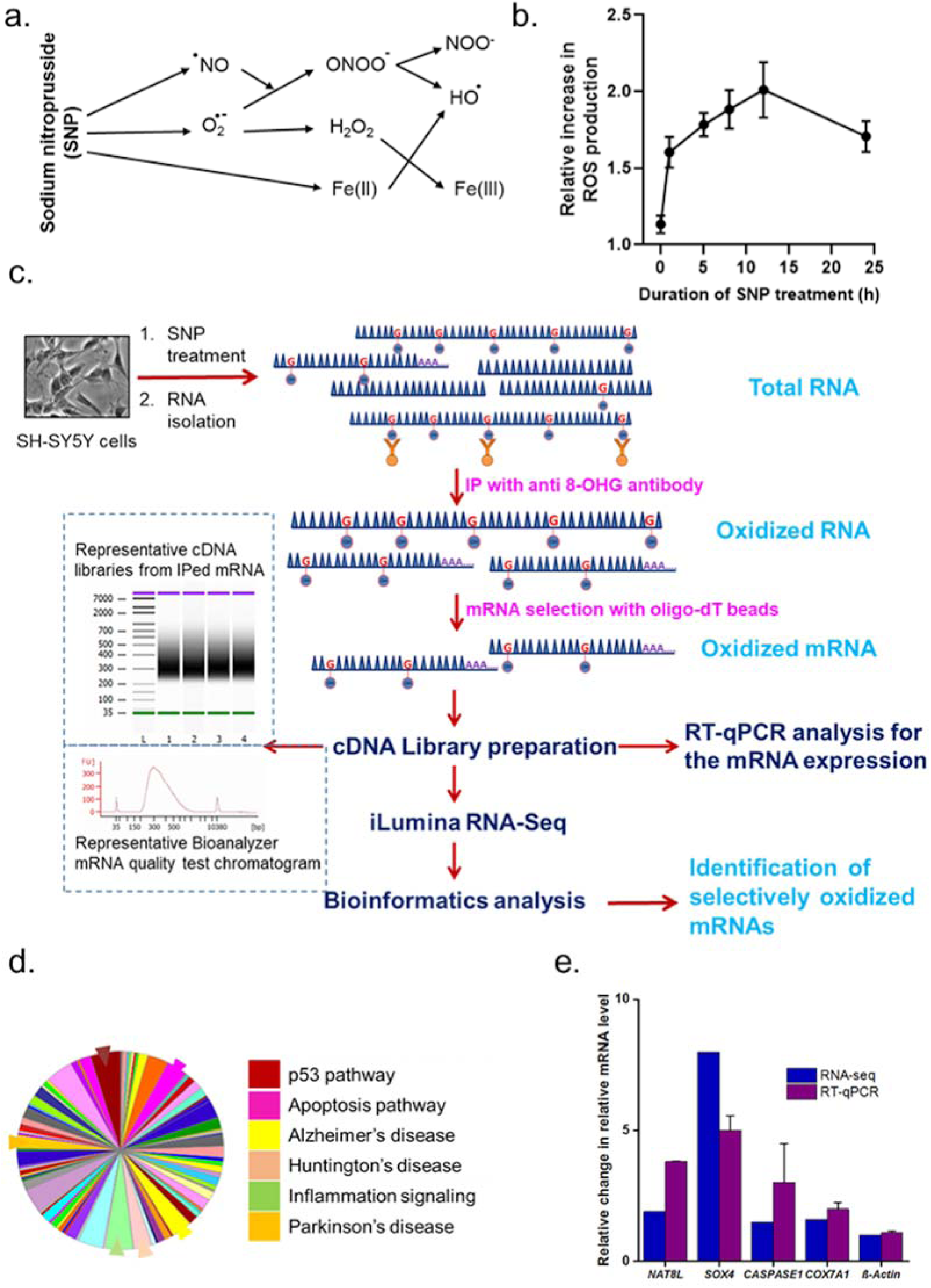
A set of the neuropathology linked mRNAs that underwent selective oxidation in SNP induced oxidative stress of the human neuronal cells. a and b. SNP induces oxidative stress in the cells by the production of ROS, c. Schematic representation of isolation, selection, quality control, and RNA-seq analysis of selectively oxidized mRNAs from the neuronal cells, d. Gene ontology analysis of the highly oxidized set of mRNA revealed that many of the neuropathologically relevant mRNAs are amongst the selectively oxidized mRNA pool, and e. RT-qPCR analyses of candidate mRNAs validate the RNA-seq data.

The RNA-seq data was analyzed as detailed in the Methods section. Overrepresentation test of the differential expression of oxidized mRNA pool was performed by **P**rotein **An**alysis **Th**rough **E**volutionary **R**elationships (PANTHER^®^) pathway analysis. PANTHER analysis identified and ranked 819 selectively oxidized transcripts contributing to various signaling pathways. The transcripts were analyzed to determine their presence in the five major categories: (i) Cellular signaling pathways, (ii) Biological process, (iii) Protein class, (iv) Cellular component, and (v) Molecular function (SI Appendix, Figure S2). Figure 1d shows the distribution by percentages of genes for various signaling pathways. Among the signaling pathways, one of the most represented class was neuropathological pathways related transcripts (represented by Alzheimer’s, Parkinson’s and Huntington diseases related pathways) and the second was inflammation-signaling pathway (Fig 1d). A third class of significantly oxidized mRNAs related to neurological disorders were the enzyme encoding mRNA transcripts, for example, *NAT8L* mRNA (SI Appendix, Figure S2). Initial analysis suggested that transcripts relevant to neuropathology were among the ones that underwent selective oxidation when human neuronal cells were subjected to mild oxidative stress.

To validate the RNA-seq data we performed RT-qPCR analyses of some of the selectively oxidized mRNAs. As illustrated in Figure 1e, many of the tested mRNAs showed a higher expression in RT-qPCR analyses of immunoprecipitated mRNA (compared between SNP IP mRNA vs control mRNA), hence validating the RNA-seq data.

Because *NAT8L* mRNA is one of the selectively oxidized candidates identified via RNA-seq analysis of oxidized RNA pool with a possible link to MS pathogenesis with key catalytic function in neurons we decided to focus our analysis on the downstream influence of *NAT8L* mRNA oxidation. NAT8L is a neuron-specific N-acetylaspartate (NAA) biosynthetic enzyme, catalyzing the NAA synthesis from L-aspartate and acetyl-CoA (Figure 2a and b). The protein sequence of NAT8L is very well conserved in mammals; with a >95% sequence conservation between mouse and human (Figure 2c). Because NAT8L directly catalyzes NAA synthesis and lower NAA level is a consistent feature in major neuropathological conditions, such as MS and Parkinson’s disease, investigating the impact of oxidation on *NAT8L* mRNA is an obvious choice (25, 30, 32, 41, 42). Although, the reduction of NAA in the MS CNS is well established, implying a crucial role of NAA deficiency in MS progression, the reason behind the reduction is not known. This led us to reason and reiterate that the selective oxidation of *NAT8L* mRNA in MS neurons could be linked to NAA deficiency in MS brain, which can affect myelin biosynthesis and contribute to MS pathology.

**Figure 2.**
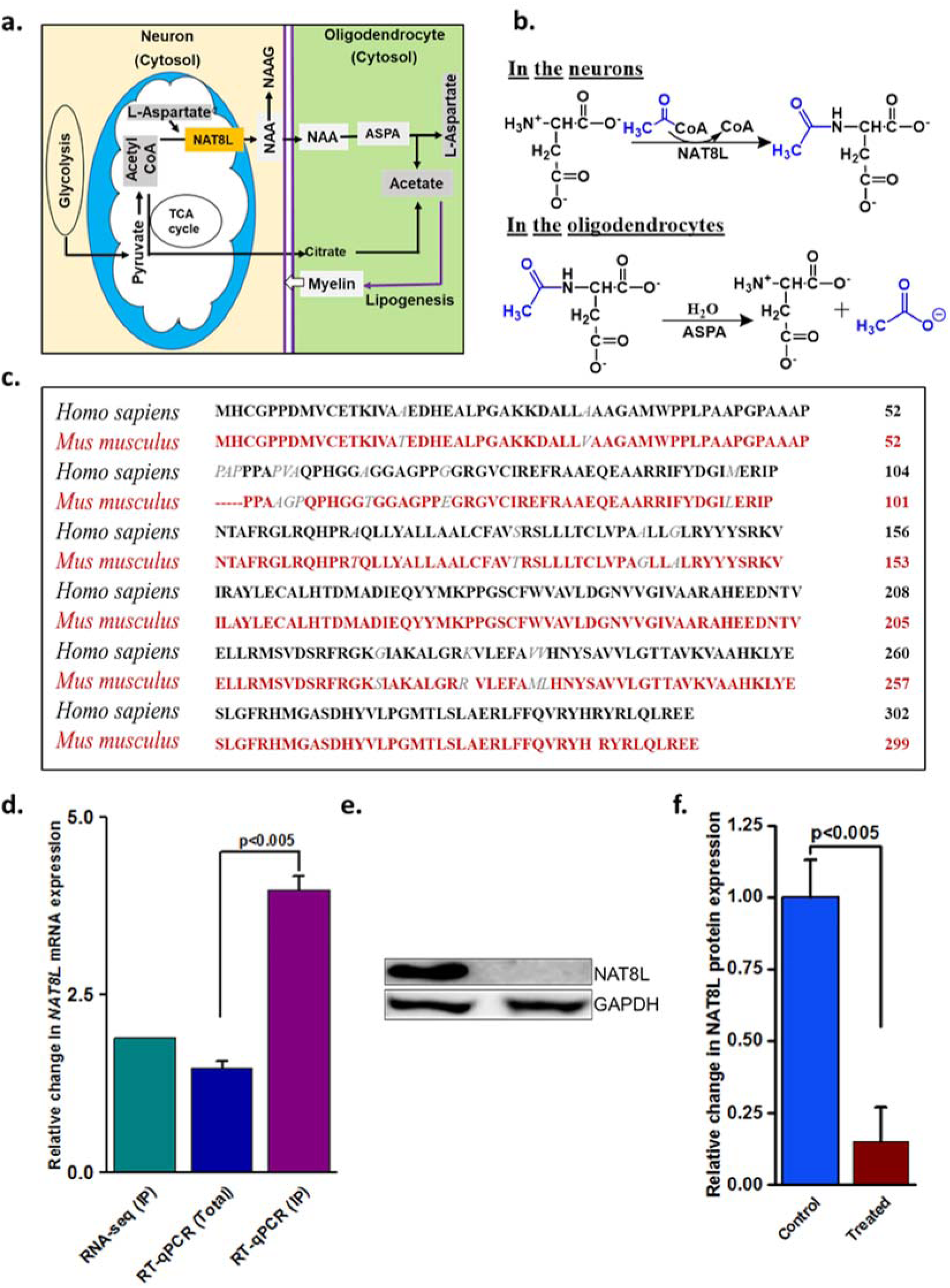
Under oxidative stress NAT8L protein expression is affected by *NAT8L* mRNA oxidation in human neurons. a. Schematic representation of the central role of NAT8L in NAA metabolism in brain, b. The catabolic and anabolic reactions of NAA, c. NAT8L sequence is highly conserved in the mammals with a 95% sequence conservation between humans and mice, d. NAT8L mRNA undergoes selective oxidation in human neurons under oxidative stress, e. A representative western blot analyses of total protein extracted from SH-SY5Y cells, and f. quantitation of three western blots as represented in e.

We observed nearly two-fold increase in *NAT8L* mRNA in the oxidized pool by RNA-seq analysis (Figure 2d) and a four-fold increase in the expression level of *NAT8L* in immunoprecipitated oxidized mRNA pool based upon the RT-qPCR data. Thus, we were able to corroborate our finding on NAT8L mRNA oxidation by two separate assays, RNA-seq and RT-qPCR. Notably, no significant change in total *NAT8L* mRNA expression was observed after SNP treatment as established by RT-qPCR (Figure 2d).

### NAT8L protein is downregulated in SNP treated human neurons

Oxidized mRNAs have been suggested to result in truncated proteins or reduced proteins or both (13, 20). Because *NAT8L* mRNA is selectively oxidized under oxidative stress in the human neuronal cells and its dysregulation is implicated in MS, we wanted to test the impact of *NAT8L* mRNA oxidation on its cognate protein expression. To test the impact of mRNA oxidation on the protein level, the cell extract from the untreated and the SNP-treated neurons were separated by SDS-PAGE followed by immunoblotting for NAT8L protein. The protein expression was normalized with the expression of GAPDH protein as the internal control. Interestingly, we observed a nine-fold reduction in the NAT8L protein level (Figure 2e and f) in SNP treated neuronal cells. This observation establishes a connection between oxidatively damaged *NAT8L* mRNA to a reduced NAT8L protein expression in human neurons under oxidative stress.

### *Nat8l* mRNA is highly oxidized in the brains of cuprizone-fed mice

Next, we embarked upon studying the relative level of *Nat8l* mRNA oxidation and its possible impact on protein expression in an MS animal model. The cuprizone-fed mouse model is one of the common toxicant induced MS animal models (43). The inclusion of copper chelator compound cuprizone in the diet is known to induce demyelination of neurons in the brain resulting in MS like phenotype in mice (43). Although, the exact mechanism of cuprizone induced demyelination is yet to be understood clearly, the increase in ROS in the brains of cuprizone-fed mice is well documented (44). This led us to rationalize that the cuprizone-fed mouse model would be a useful tool to study the effect of OS in MS in combination with the comparison to the *in cellulo* data. The mice were fed cuprizone containing diet for six weeks (regular diet for control mice) and were sacrificed according to an already established protocol, after which we harvested the brains for downstream analyses.

Initially, we had to establish if RNA in the brain of cuprizone-fed mice undergo oxidation. For that purpose, we isolated the total RNA from the gray areas of the cuprizone-fed mouse brain and the control mouse brain and immunoprecipitated both groups with anti-8-OG antibody. We quantitated the amount of oxidized RNA in both pools and found a 2-fold increase in total RNA oxidation in cuprizone-fed mouse brain in comparison to that from the control mouse brain (Figure 3a). Next, we analyzed the relative change in *Nat8l* mRNA level in cuprizone-fed mouse brain. RT-qPCR analysis of total mRNA pool from both groups of mice did not show a significant change in *Nat8l* mRNA expression (Figure 3b). However, while analyzing the expression in immunoprecipitated oxidized mRNA, we observed a 2-fold more *Nat8l* mRNA in oxidized mRNA pool of cuprizone mouse brain compared to the total RNA in control mouse brain or cuprizone-fed mouse brain (Figure 3b). This is exactly the same trend we observed in human neuronal cells under SNP treatment (Figure 2). Thus, there is a direct correlation between the oxidative damage of *NAT8L* mRNA *in cellulo* and in the brain of the MS mouse model.

**Figure 3.**
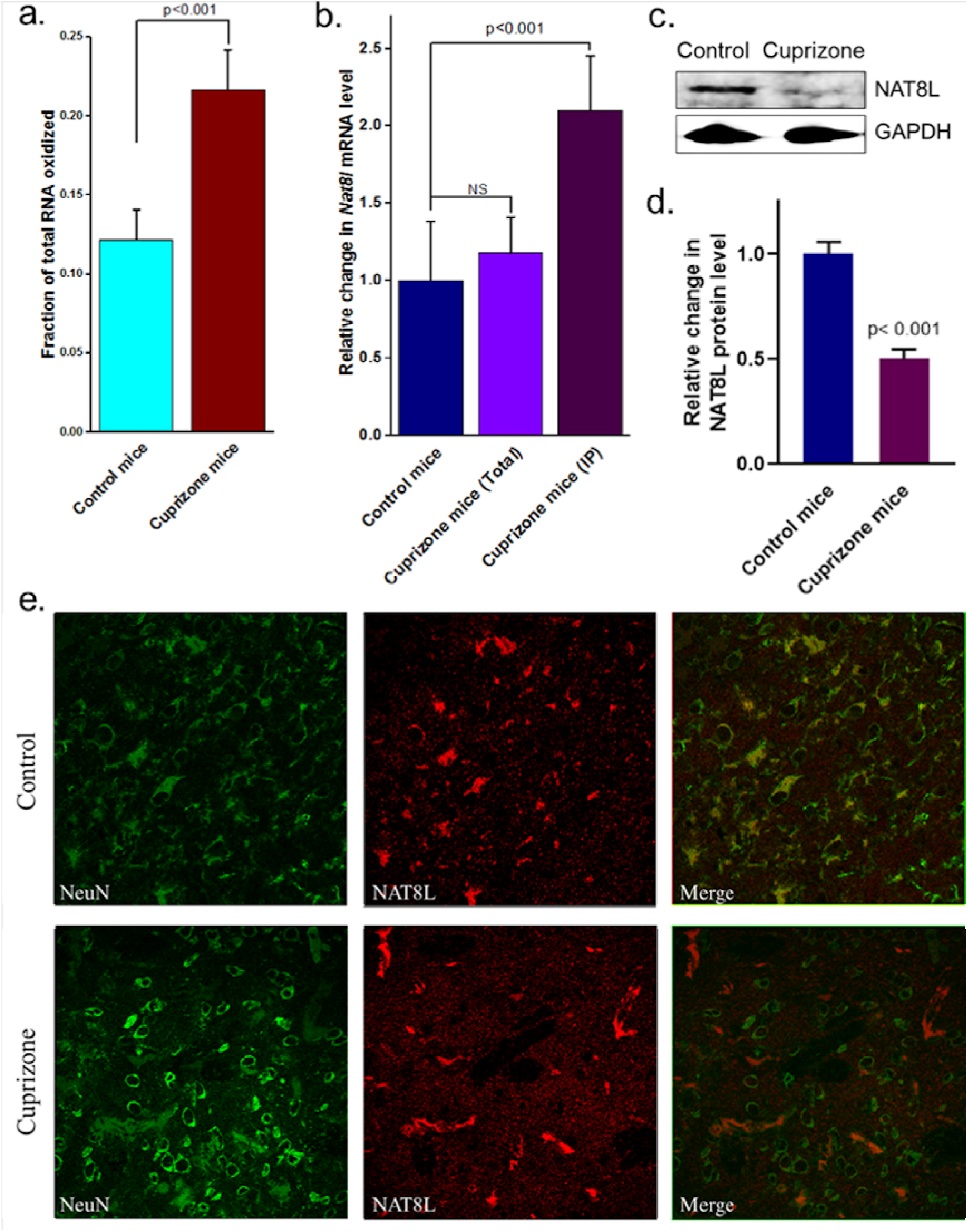
*NAT8L* mRNA oxidation is linked to a reduced NAT8L protein production in cuprizone-fed mouse brain. a. Immunoprecipitation of total RNA with 8-OG antibody from the gray areas of mouse brain followed by spectrophotometric quantification revealed two-fold more RNA oxidation in cuprizone-fed mouse brain, b. *Nat8l* mRNA is more oxidized in cuprizone-fed mouse brain compared to the mouse fed with normal diet, n=3, statistical significance was calculated using ordinary one-way ANOVA, multiple comparisons with Tukey’s multiple comparisons test (Graphpad Prism 8), c. NAT8L protein expression is compromised in cuprizone-fed mouse brain, a representative western blot analyses of total protein extracted from the gray areas of mouse brain, d. quantitation of three western blots as represented in c, and e. Immunohistochemical analysis of 30 µm thick mouse brain sections with anti-NAT8L antibody supported the reduction in NAT8L protein expression in within the neurons (anti-NeuN antibody, green channel).

### NAT8L protein is downregulated in the brain of cuprizone-fed mouse

Cuprizone-fed mice are characterized by a severe demyelination of neurons (43). A reduced availability of NAA could be a contributing factor to this demyelination process since NAA in the oligodendrocytes is known to contribute in myelination process by providing acetate needed for lipogenesis (29, 30). *In cellulo* studies showed that the oxidatively damaged *NAT8L* mRNA lowers NAT8L protein expression, in turn reducing NAA level, which prompted us to investigate whether NAT8L protein expression is influenced similarly (as in case of SNP treated neurons) by the oxidation of *Nat8l* mRNA *in vivo*. Total protein was isolated from the gray matter of control and cuprizone mice brains, separated in SDS-PAGE and immunoblotted against anti-NAT8L antibody. Remarkably, we observed a 50% reduction in the NAT8L protein level in cuprizone-fed mouse brain compared to the control (Figure 3c and d).

In a separate experiment, we performed immunohistochemical analyses of brain tissue sections to analyze the change in NAT8L level in cuprizone-fed mouse brains. As demonstrated in Figure 3e, our findings showed a sharp reduction to no expression in the overlay of NAT8L staining and NeuN (neuronal marker protein) staining, clearly indicating the reduction in NAT8L protein in the neurons of cuprizone-fed mouse brain. We also wanted to test whether NAT8L is expressed in non-neuronal cells. Co-immunostaining of NAT8L protein with GFAP (Glial fibrillary acidic protein, astrocyte marker) and MBP (Myelin basic protein, oligodendrocyte marker) proteins showed there is no significant overlap of NAT8L antibody in the non-neuronal cells implying an exclusive neuronal presence of NAT8L protein (SI Appendix, Figure S3) under the tested conditions.

### Under oxidative stress enzymatic activity of NAT8L is reduced in human neurons

N-Acetyl aspartate transferase 8 like protein catalyzes the transfer of an acetyl group from acetyl-CoA to aspartate resulting in NAA (Figure 2 and 4) in mammalian neurons. It has been shown that the membrane bound NAT8L and other NAT proteins preserve their catalytic activity in the total cell extract (45). Our goal was to analyze the impact of reduction in NAT8L (belongs to Asp-Nat family of enzymes) protein level on its enzymatic activity in the total cell extract of neuronal cells and in the extract from cuprizone-fed mouse brain. This analysis would help us answer two crucial questions: 1) whether the observation of a significant reduction in NAT8L protein, as measured via western blot *in cellulo* and *in vivo* extracts is biologically significant and 2) is there an alternative Asp-Nat mechanism in the absence of NAT8L protein in mammalian neurons? We performed the acetyl group transfer reactions from acetyl coenzyme A to ^14^C labelled aspartic acid as detailed in the methods section. We observed a reduction in enzymatic activity in cell extracts from the SNP treated neurons in comparison to the control cells (Figure 4a and b). We also observed a similar trend in the Asp-Nat enzymatic activity in the extract from cuprizone-fed mouse brain (SI Appendix, Figure S4). These data indicate that the reduction in NAT8L protein is detrimental to NAA synthesis in SH-SY5Y cells as well as in mouse brain and it appears that there is no compensatory alternate mechanism that can synthesize NAA in the neurons when NAT8L protein synthesis is compromised.

**Figure 4.**
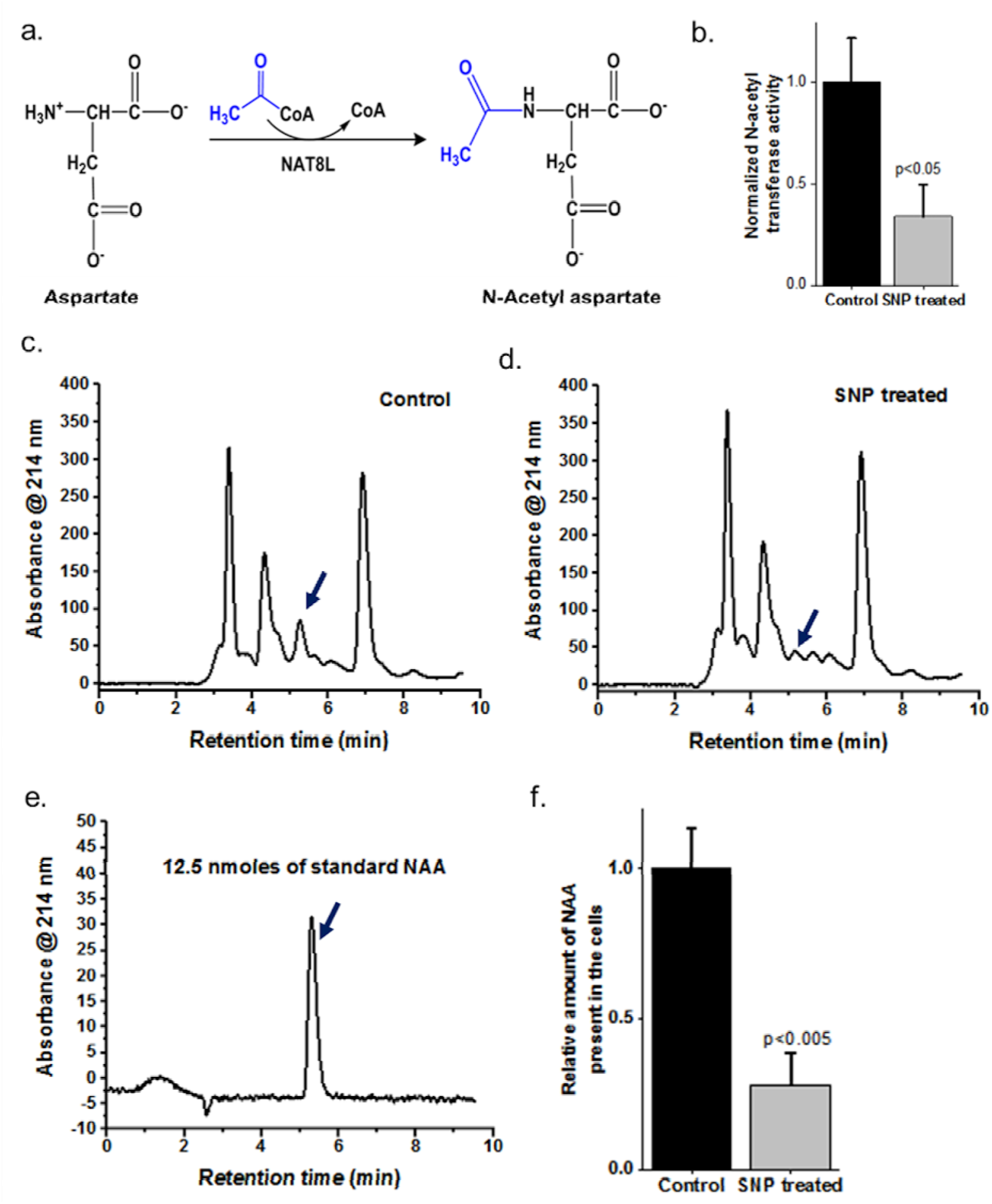
Aspartyl N-acetyl transferase (Asp-Nat) activity is compromised in the SNP treated neuronal cells. a. the reaction catalyzed by NAT8L in the neurons, b. NAT8L enzymatic activity is reduced in SNP treated neurons as measured by the quantitation of the transfer of acetyl group to the radiolabeled aspartate, c & d. HPLC chromatograms showing a reduction in NAA level in SNP treated neurons in comparison to the non-treated neurons (blue arrows indicate the peaks corresponding to NAA), e. A representative HPLC chromatogram of standard NAA, and f. Normalized representation of quantitation of the change in NAA level between SNP treated and non-treated neuronal cells from three independent experiments.

### NAA level is reduced in SNP treated neuronal cells

If NAT8L is a key enzyme that is responsible for the synthesis of NAA in human neurons, the reduced expression of this protein should also result in the reduction of NAA level. To test this hypothesis, we quantitated the amount of NAA in the control versus SNP treated SH-SY5Y cells by HPLC (Figure 4 c-f). Cell numbers were normalized after SNP treatment and before NAA isolation. Our data demonstrate that after 8 h of treatment with SNP the NAA level in SH-SY5Y cells was reduced by about 75%. The reduction in the amount of the crucial neuronal metabolite NAA in the neuronal cells corroborates well with a higher level of *NAT8L* mRNA oxidation and subsequent reduction in NAT8L protein level as was observed via immunoblot analysis.

### NAT8L protein is significantly reduced in the normal appearing areas of MS brain and is reduced further in the MS lesions

There are reports on the reduction of NAA level in the MS brain (32, 35, 46). In fact, the reduction of NAA in the brain has been used as a marker for detecting mitochondrial dysfunction in several neurological disorders including MS (32, 35). In spite of being an indicator of mitochondrial dysfunction and poor neuronal health, the cause of reduction in NAA is poorly understood. The observation of reduced NAT8L protein and reduced Asp-Nat activity in the oxidatively stressed neurons and MS mouse model prompted us to measure the level of oxidation in *NAT8L* mRNA and change in NAT8L protein level in normal appearing gray matter (NAGM) of postmortem MS brain. RT-qPCR analysis of immunoprecipitated mRNA isolated from NAGM of postmortem MS brain showed a higher oxidation of *NAT8L* mRNA (Figure 5a). Next, we investigated whether the impact of oxidatively damaged mRNA is reflected in the level of protein synthesized. The immunoblot analyses of the protein extracted from the normal appearing gray matter in MS and non-MS brain revealed that there is indeed about 40% reduction in the relative amount of NAT8L protein in the normal appearing gray areas of MS brain compared to a similar area of the non-MS brain (Figure 5 b and c). Because we observed a sharper NAT8L decline in cuprizone-fed mouse gray matter, which is more relevant to demyelinated MS lesions, we further wanted to analyze the presence of NAT8L in lesioned areas of postmortem MS brain. To our no surprise, we observed nearly 85% reduction in NAT8L protein in MS lesions (Figure 5 d-f).

**Figure 5.**
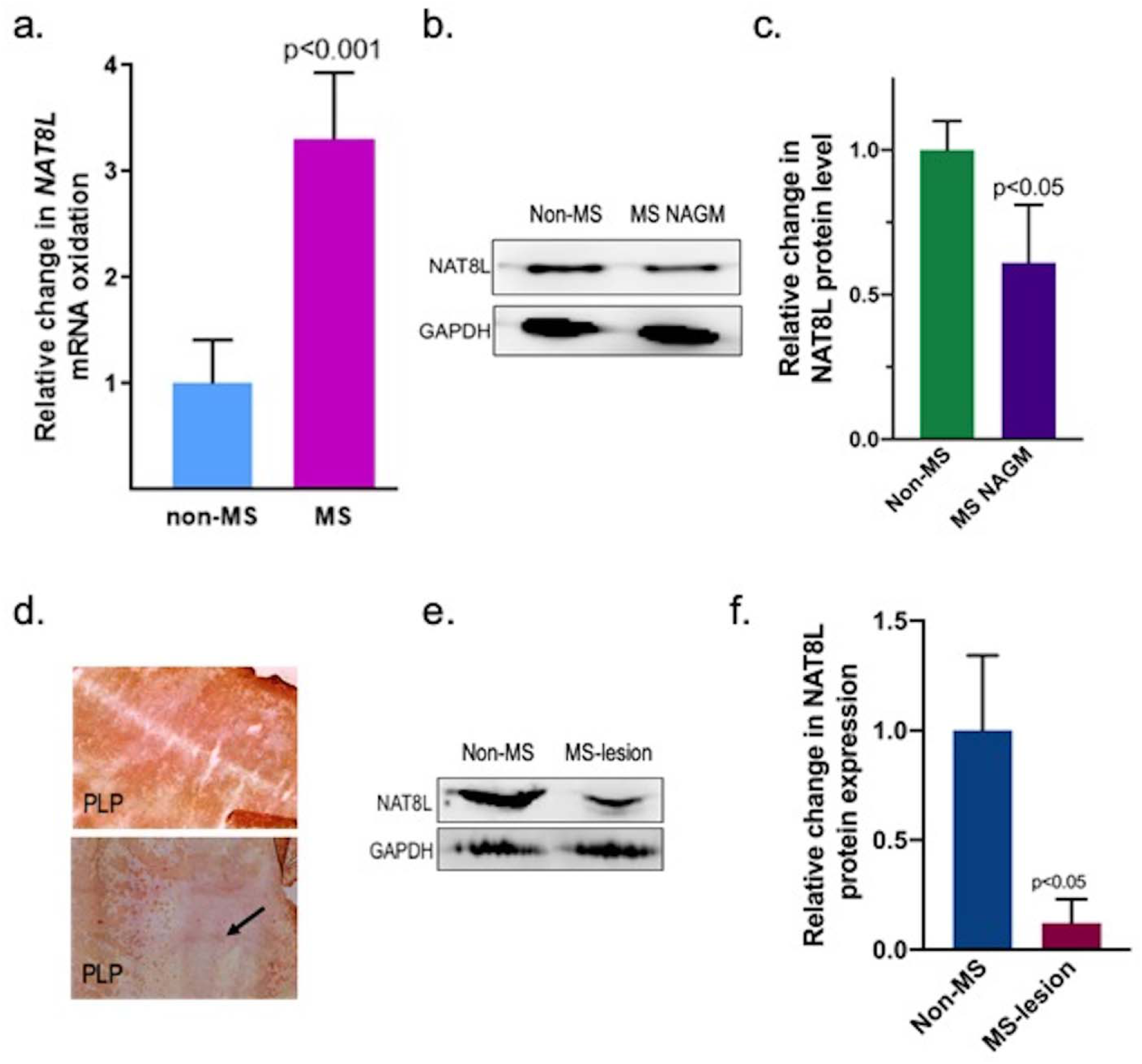
Messenger RNA oxidation in MS brain is linked to reduced NAT8L protein production. a. RT-qPCR analyses of 8-OG immunoprecipitated RNA from the non-MS and MS brains show a three-fold higher oxidation in NAT8L mRNA in the normal appearing gray matter (NAGM) of MS brain, b. Western blot detection of NAT8L protein from the extract isolated from NAGM of postmortem MS brain and the gray matter of non-MS brains, c. Quantification of three independent western blots from b, d. PLP stain of MS brain sections to demonstrate normal appearing (top) and lesion containing (bottom) areas, e. Western blot demonstrating the reduction of NAT8L level in MS lesions, and e. Quantification of three independent western blots from e.

Thus far, we have shown that under oxidative stress *NAT8L* mRNA is oxidized in human neuronal cells, cuprizone-fed mouse brain, and in post-mortem MS brain resulting in reduced NAT8L protein production and lowered NAA level. The lower NAA level in the normal appearing gray matter of postmortem MS brain and cuprizone-fed mice was reported previously (35, 44). Therefore, our data provide a molecular rationale for the decreased NAA level in MS brain.

### *In vitro* translation of oxidized *NAT8L* mRNA show a reduced protein production

To support the *in cellulo* and *in vivo* data on the oxidation of *NAT8L* mRNA, the reduced level of NAT8L protein and a lowered NAA level respectively, we designed a direct assay to demonstrate the effect of *NAT8L* mRNA oxidation on its translation. For that purpose, a commercially available *in vitro* translation system was used. *NAT8L* mRNA was transcribed from the pPB-C-His-NAT8L vector and the transcribed mRNA was oxidized using Fenton reaction conditions as described in the Methods section. The oxidized mRNA thus obtained was purified, quantified, and analyzed to determine the extent of oxidation. The amount of oxidized mRNA was normalized to non-oxidized mRNA before separately translating them *in vitro*. We observed a 70% reduction in the amount of full length translated product in the oxidized mRNA compared to the protein from the non-oxidized mRNA (Figure 6). This observation directly established the detrimental effect of *NAT8L* mRNA oxidation on its cognate protein synthesis.

**Figure 6.**
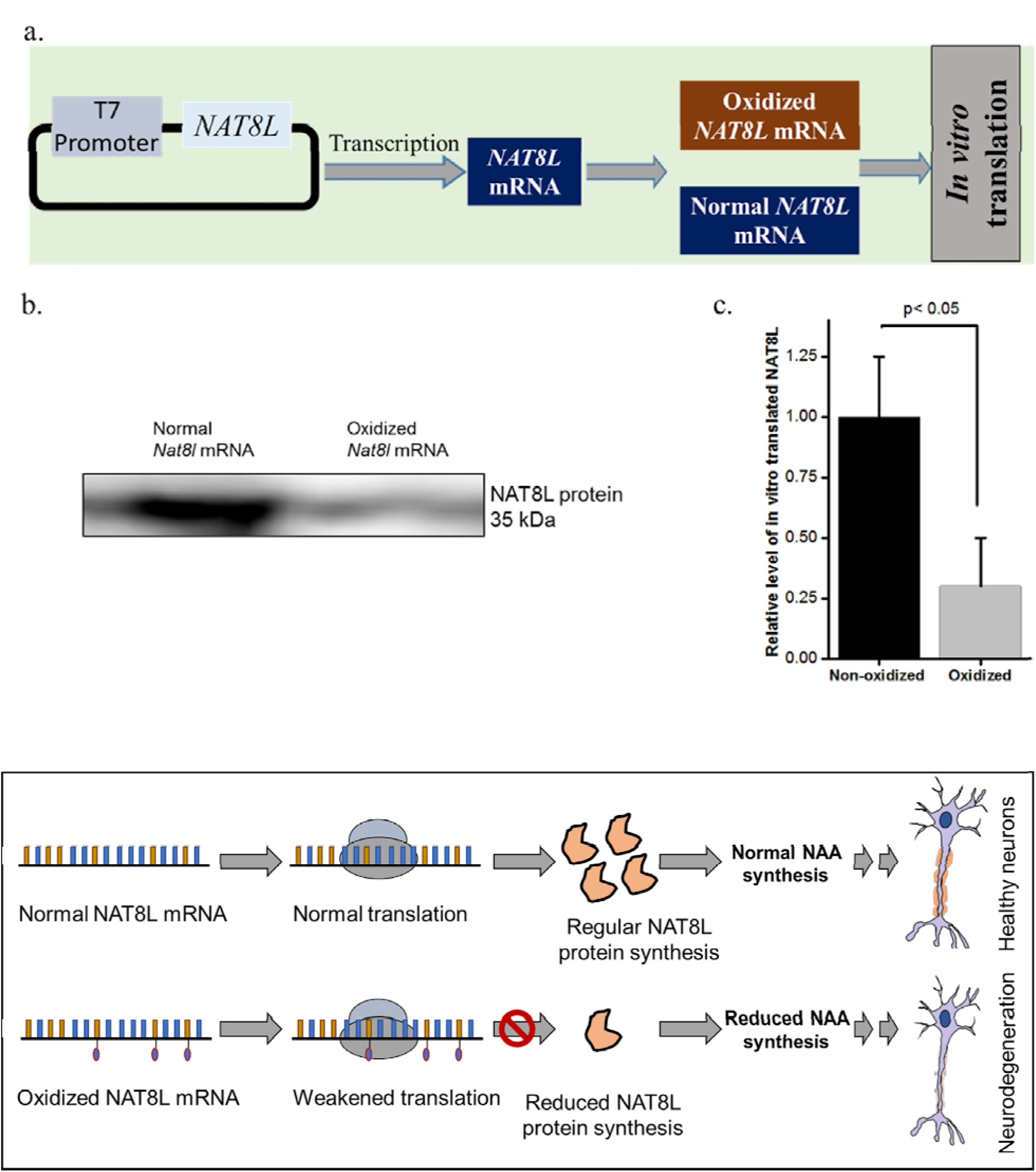
*NAT8L* mRNA oxidation inhibits its translation *in vitro*. a. Schematic of the experimental design- the *NAT8L* mRNA was transcribed from an engineered plasmid vector. The transcribed mRNA (with or without mRNA oxidation) was translated using rabbit reticulocytes-based translation system and b and c. Western blot detection of translation product using anti-NAT8L antibody and their quantification.

**Figure 7.** A proposed model to demonstrate the role of *NAT8L* mRNA oxidation in poor NAA synthesis, demyelination of axons, and MS neurodegeneration.

## Discussion

In this report we show that oxidatively damaged *NAT8L* mRNA results in reduced NAT8L protein production and concomitant reduction in the level of NAA, linking mRNA oxidation to neurodegeneration in MS. Initially we established that there is mRNA oxidation in human neuronal cells under OS, cuprizone-fed mouse brain and post-mortem MS brain tissue, all linking mRNA oxidation to MS. Previous studies have indicated that RNA oxidation is an early event in the pathological cascade of AD neurodegeneration (11, 16). Increased RNA oxidation has been observed in post-mortem brains of subjects with mild cognitive impairment (47) and pre-symptomatic subjects with a familial AD mutation (10). Studies on AD revealed that neuronal RNA oxidation precedes the formation of amyloid β or tau pathology, the later are implicated in AD neurodegeneration (48). Increased RNA oxidation was also observed in the pre-symptomatic stage in transgenic mice expressing an ALS-linked mutant SOD1^G93A^, suggesting that RNA oxidation is a common early event preceding motor neuron degeneration in ALS (13). Thus, our data on selective mRNA oxidation in MS and MS models are consistent with studies in neurodegenerative diseases, such as, AD and ALS.

Among the pool of total mRNA analyzed from the oxidatively stressed human neuronal cells, only 819 were significantly expressed in the oxidized mRNA pool. Thus, our observation that only a subset of mRNAs undergo detectable oxidation is consistent with the previous report by Kong, et al (16). The authors reasoned that the selective mRNA oxidation is neither due to the abundance of mRNA species nor due to any common structural feature that could explain the selectivity (12, 13). We discovered that many of the neuropathologically relevant transcripts are amongst the selectively oxidized group of mRNAs. The pathway analysis of the highly oxidized transcripts showed that many of them are members of inflammatory pathways and/or AD, PD, and HD related pathways, all of which could be relevant to MS neurodegeneration (Figure 1d). While trying to decipher the reason behind the selectivity in mRNA oxidation, we asked two question: 1) whether a high G-density in the transcripts corresponds to a higher mRNA oxidation, given the increased susceptibility of Gs to undergo oxidation, and 2) whether the organelle distribution of respective proteins contribute to the selectivity in mRNA oxidation? In our study, neither of these factors seemed to contribute towards the selectivity of mRNA oxidation. Among the 819 oxidized mRNAs, some contain as low as 17% Gs in the transcript, while others contain as high as 44% Gs (SI Appendix, Figure S5). Similarly, some of those transcripts code for mitochondrial proteins while some others code for cytoskeletal and nuclear proteins (SI Appendix, Figure S2).

The occurrence of detected oxidized mRNAs in various pathways linked to neurodegenerative diseases and inflammation provided us with a platform to undertake detailed mechanistic studies on the impact of individual mRNA oxidation to the neuronal health under oxidative stress. Based on the growing interest in NAT8L protein for its link to brain physiology, and recent reports from the McDonough lab that NAA deficiency and MS pathology are linked (32, 35), *NAT8L* was chosen as a candidate mRNA from the oxidized mRNA pool for further analyses. *NAT8L* mRNA is one of the selectively oxidized mRNAs under the MS microenvironment and its cognate protein does have an enzymatic activity that is responsible for the synthesis of the metabolite NAA, which is a key metabolite contributing to the synthesis of healthy myelin sheath (45). Because the oxidized mRNA is known to undergo reduced translation (20), we expected to observe a reduced level of NAT8L protein due to the oxidative damage of its cognate mRNA. Indeed, we consistently observed reduction in NAT8L proteins *in cellulo*, in cuprizone-fed mouse model and in the post-mortem MS brain. Additionally, we observed a reduced NAT8L protein production from the translation of *in vitro* synthesized and oxidized *NAT8L* mRNA, clearly demonstrating that the oxidation of *NAT8l* mRNA directly causes reduction in the NAT8L protein. The reduction in the NAT8L protein level in cuprizone-fed mouse brain corroborates well with previous findings on the decrease in NAA level in cuprizone-fed mouse brain (44). Analysis of Asp-Nat activity showed a reduction in the NAT8L enzymatic activity in the extract from oxidatively stressed neuronal cells and in the extract from the cuprizone-fed mouse brain. In both situations, the loss in Asp-Nat activity proved to be a key in the reduction of NAA. As we discovered, higher mRNA oxidation and the reduction of NAT8L protein in normal appearing gray areas of postmortem MS brain seems to be related with the reduced NAA level and demyelination in MS brain. These findings correlate well with the reduced level of NAA reported in post-mortem MS brain (32, 33, 35). Taken together, these results clearly establish a link between higher ROS production, selective oxidation of *NAT8L* mRNA and the detrimental impact it has on its own translation both *in cellulo* and in an MS animal model resulting in reduced NAA level.

The *NAT8L* mRNA contains several G-rich stretches that can potentially adopt intramolecular G-quadruplex (GQ) structures. There are multiple G-stretches embedded within the ORF that can potentially form GQ structures. Interestingly, the deletion within this G-rich sequence in *NAT8L* ORF has been shown to result in highly reduced NAA level in the only case reported in a patient with hypoacetylaspartia (45). Since the G-quadruplex (GQ) is destabilized when G’s are oxidized (49), it could be possible that a disrupted GQ in the coding region could be detrimental for the stability of the resultant protein. In addition to the ORF, the *NAT8L* mRNA sequence has at least a dozen potentially GQ forming sequences (PGQSs) in the 3’-UTR. The GQs in the 3’-UTR of mRNAs have been suggested to play roles in mRNA stability (50). Thus, oxidation of G’s in the *NAT8L* mRNA 3’-UTR could result in destabilization of one or more of the PGQSs in turn causing mRNA instability. It can potentially affect the mRNA surveillance mechanism by disrupting RNP complexes designed to store translating machinery under stress.

Given the main pathology of MS is defined by demyelination, a reduced NAA level will be detrimental to compact myelin synthesis. In this work we showed that NAT8L mRNA oxidation results in reduced NAT8L protein leading to decreased NAA level. The McDonough lab has previously showed that NAA is required for a proper myelin structure as found by a less compact myelination in *Nat8l*^*-/-*^ mouse (32). Thus, we can connect NAT8L mRNA oxidation to reduced or weak myelin synthesis and neurodegeneration in MS. This finding can also be relevant in other demyelinating diseases, such as, Balo’s disease and Devic’s Disease (51). Our discovery about the direct link between selective *NAT8L* mRNA oxidation to the low abundance of the critical neuronal metabolite NAA could also inspire other possible directions in MS and neurological disorder research. For example, NAA in the brain is not only related to a healthy myelin production but also acts as a player in mitochondrial health (25). A reduced NAA level in neurons is a marker of mitochondrial dysfunction. NAA is the precursor of NAAG in the neurons, which is another important signaling molecule that plays a role in astrocytes where it is known to be involved in pain signaling via glutamate receptor mediated pathway (24, 46, 52). A reduced expression of NAT8L enzyme in neurons could result in reduced production of NAAG in neurons thereby altering the pain signaling pathway.

Overall, selective oxidation of *NAT8L* mRNA in MS neurons, the reduced amount of NAT8L protein expressed under such an environment and our data on *in vitro* translation of oxidized *NAT8L* mRNA show a direct inverse correlation between mRNA oxidation and NAT8L protein expression. We thus conclude that ROS induced oxidation of *NAT8L* mRNA results in the reduced production of NAT8L protein in MS neurons hence debilitating NAA metabolism (Figure 7). To the best of our knowledge, our findings begin to address the direct pathological connection between selective mRNA oxidative damage and MS neurodegeneration. We expect that this work will provide a roadmap for analysis of the other neurological disorders that are associated with mRNA oxidation.

## Methods and methodology

### Cell culture, SNP treatment, and ROS measurement

Human neuroblastoma SH-SY5Y cells obtained from ATCC were cultured in Dulbecco’s modified Eagle’s medium supplemented with 10% (v/v) fetal bovine serum, and 1% penicillin/streptomycin at 37 °C under 5% CO_2_ in air. The cells were seeded in a 6-well plate (500K cells/well) and they were allowed to settle for 24 h before the treatment. To determine the effect of SNP on retinoic acid differentiated SH-SY5Y cells, the cells were treated with various doses of SNP for 8 h. In a single experiment, each treatment was performed in triplicate. The production of ROS in the cells after SNP treatment was measured using ROS-Glow™ H_2_O_2_ assay following the manufacturer’s protocol (Promega).

### Brain extract preparation

MS and non-MS postmortem frozen brain tissue blocks were obtained from the Rocky Mountain MS Center and the Brain and Spinal Cord Resource Center at UCLA under Institutional Review Board guidelines. MS and non-MS tissue were matched for brain region, age, and postmortem interval as closely as possible (SI Appendix, Table S1). Respective tissue blocks were used to cut 30 um tissue sections to evaluate for normal appearing vs lesion containing areas with PLP staining. RNA and protein were extracted from normal appearing gray matter and lesion containing gray matter of the MS postmortem brains and gray matter of the non-MS brains.

### RNA isolation and immunoprecipitation

RNA was isolated from SH-SY5Y cells using standard protocol (TRIzol™ Reagent, Invitrogen) and immunoprecipitated with anti 8-OG antibody (QED Biosciences) as described before (6). 25 μg of total RNA was incubated with 30 μg of anti 8-OG antibody at room temperature for 2 h. 35 μL of immobilized protein L agarose gel beads (Pierce) were added to the RNA-antibody mixture and incubated overnight at 4 °C. The beads were washed three times (3 × 5 min) with 200 μL PBS with 0.04% (v/v) Nonidet P-40 (Roche Applied Science). The oxidized RNA: antibody: protein L agarose beads complexes were separated from non-oxidized RNAs (which remained in the supernatant) by centrifugation at 1500 rpm for 5 min at 4 °C. Oxidized RNA-bead complexes were mixed with following reagents: 3 mL of PBS with 0.04% Nonidet P-40, 300 μL of 10% (w/v) sodium dodecyl sulfate (SDS), and 3 mL of PCI (phenol: chloroform: isoamyl alcohol; 25:24:1), and the mixture was incubated at 37 °C for 30 min (with occasional vortexing) and separated to an aqueous and an organic phase by spinning at 10,000 rpm for 15 min at 4 °C. The aqueous layer containing oxidized RNA was separated and mixed with 40 μL of 3 M sodium acetate buffer (pH 5.3), 2 μL of 10 μg/μL glycogen and 1 mL of absolute ethanol. The sample was then frozen at −80 °C for 1 h and centrifuged for 20 min at 13,200 rpm at 4 °C. The pellet was washed with 70% ethanol, vacuum dried and dissolved in T_10_E_0.1_ buffer. Oligo d(T)_25_ magnetic beads were used to enrich oxidized mRNA using the manufacturer’s protocol (New England Biolabs).

RNA from mouse brains (control and cuprizone-fed) and postmortem human brain sections (MS and non-MS) was isolated from the homogenized gray matters following the same protocol.

### RNA sequencing and Bioinformatics

Messenger RNA from SH-SY5Y cells (oxidized RNA from SNP treated cells and total mRNA from non-treated cells) was obtained as described above. The mRNA was tested for their quality using Agilent Bioanalyzer test and the cDNA libraries were synthesized using iLumina Trueseq library preparation kit. cDNA libraries were sequenced using iLumina HighSeq platform (75 base pair sequencing, 40 million reads per sample, girihlet Inc, https://www.girihlet.com/). RNA-Seq data were analyzed using CLC Genomics Workbench software v12.0 (Qiagen Digital Insights)- a software suite that provides a variety of tools for next-generation sequencing analysis. The RNA-seq workflow, as described in the CLS Genomics manual, was followed (http://resources.qiagenbioinformatics.com/manuals/clcgenomicsworkbench/current/index.php?manual=RNA_seq_Small_RNA_analysis.html, accessed March 6, 2020). Briefly, single-end RNA-Seq reads, obtained in FASTQ format, were subjected to sequence quality controls using “QC for Sequencing Reads” available under the toolbox item “Prepare Sequencing Data”. Reads with Phred quality score > 20 were then aligned to the human reference genome GRCh38 assembly sequence using the default parameters provided by the “RNA-seq Analysis” tool available under the “RNA-Seq Analysis” toolbox item. Genes and transcripts annotations provided by Ensembl (release 92) were used. Around 94% of reads from each sample were mapped into the reference genome.

Differentially expressed genes (DEGs) between cells with and without SNP treatment were then determined using the “Differential Expressions in Two Groups” tool, also available under the toolbox item “RNA-Seq Analysis.” It uses a multifactorial statistics based on a negative binomial Generalized Linear Model. Genes with FDR corrected P-value <= 0.05 and fold change >=1.5 OR <= −1.5 were considered as differentially expressed. The expression of 1627 genes were significantly different in SNP treated (oxidized) mRNA samples as compared to untreated control. Among these, 819 genes were upregulated indicating relatively higher oxidation. A spreadsheet with the results of DEGs along with gene expression counts of individual genes for each sample was generated using the tool “Create Expression Browser” also available under the toolbox “RNA-Seq Analysis.” The spreadsheet is attached as a supplementary file (Supplementary table S3). The RNA-seq data were deposited into the Gene Expression Omnibus database (GSE144294). Overrepresentation test of highly oxidized mRNA pool (upregulated genes from RNA-seq data) was performed by Protein Analysis Through Evolutionary Relationships (PANTHER^®^) pathway analysis.

### Cuprizone mouse model

The mode of action of cuprizone-toxicity is believed to take place via the copper-chelation action of cuprizone, which inhibits mitochondrial enzymes of the respiratory chain that require copper as co-factor. This leads to oxidative stress resulting in primary oligodendrocyte apoptosis which is closely followed by microglia and astrocyte activation. These events lead to demyelination and neurodegeneration. For *in vivo* experiments, we used C57Bl6/J male mice (7 weeks of age). One group of mice were treated with regular diet and other with 0.3% cuprizone-containing diet (TD. 140805, Teklad Global) for 6 weeks. We perfused the mice with saline water and isolated the brains from mice. Isolated brains were fixed in 4% paraformaldehyde and cryoprotected with sucrose. Brains were sliced coronally using a cryostat machine. All experiments were performed according to protocol approved by Institutional Animal Care and Use Committee of Kent State University.

### Immunohistochemistry

Immunofluorescence staining: 30 μm thick tissue sections were fixed with 4% formaldehyde in PBS (pH 7.4) for 20□min, washed with PBS, and then incubated with 0.25% Triton X-100 in PBS for 15□min. The sections were treated with 10□mM citrate solution (pH 6.0) for antigen retrieval at 95□°C for 15□min, washed with PBS, and then blocked in 5% normal donkey serum (Sigma-Aldrich, St. Louis, MO) in PBS for 1□h. Next, the sections were incubated in primary antibody (anti-NeuN, anti NAT8L, anti-GFAP, anti-MBP) with 5% normal donkey serum in PBS overnight at 4□°C, washed with PBS-T, and then incubated with fluor secondary antibodies (with or without DAPI) at room temperature for 2□h. After being washed with PBS, the sections were mounted using Vectashield mounting media (Vector Labs) and imaged using Olympus 1000X confocal microscope.

DAB staining: The tissue sections were probed with anti-PLP primary antibody followed by DAB staining to distinguish lesion vs non-lesion areas in MS brain. 30 μm tissue sections were fixed in 70% EtOH at room temp for 30 min, washed, and treated with 1% H_2_O_2_ (Santa Cruz Biotechnology sc-203336A) for 30 min and rinsed with PBS. Next, the sections were blocked in 3% normal donkey serum in PBS for 30 min followed by incubation in 1:200 PLP (EMD Millipore, MAB388-100UG) with 3% normal donkey serum, 0.5% Triton-X-100 in PBS overnight at 4 °C. The washed tissue sections were incubated in 1:500 biotinylated anti mouse secondary antibody (Vector Laboratories, Inc. BA-2000) in 3% normal donkey serum, and 0.5% Triton-X-100 in PBS at room temp for 2 hours. Next, the sections were incubated in Vectastain ABC Kit (Vector Laboratories, Inc. PK-6200) for 1 h at room temp (1 drop of A, 1 drop of B in 5 mL PBS with 0.5% Triton-X-100). The sections were then incubated with DAB Peroxidase Substrate Kit (Vector Laboratories, Inc. SK-4100) (1 drop DAB buffer, 2 drops DAB reagent, 1 drop H_2_O_2_ in 5 mL aqueous solution). The sections were mounted in glycerol, coverslipped, and imaged. The images were processed using ImageJ^®^.

### RT-qPCR

Relative mRNA levels were quantitated by RT-qPCR using mRNAs isolated from SH-SY5Y cells, gray matters of cuprizone-fed mice brains, and post-mortem MS brain. RT-qPCR was performed using SYBR Green (Quanta Biosciences) master mix reagent and gene-specific primers. Relative levels of mRNA expression in control and treated cells and control and cuprizone mice were determined by the 2^−ΔΔCt^ method. The primers used for RT-qPCR were purchased from Integrated DNA Technologies (IDT) and are listed in the supplementary information (SI Appendix, Table S2).

### Western blotting

Protein lysate (Cell lysate, or tissue homogenate) was run in a SDS polyacrylamide gel (12% separating gel and a 5% stacking gel) at 180V. Separated protein in the gel was transferred to Amersham Protran Premium 0.45 μm Nitrocellulous membrane (GE Healthcare) at 30V for 20 hours at 4 °C. The blot was blocked with 5% skimmed milk in 1X TBST buffer for 45 minutes at room temperature followed by incubation in primary antibodies (1:500, Nat8l antibody, Invitrogen PA5-68424, or 1:5000 GAPDH antibody, Santa Cruz Biotechnology, sc-365062) overnight at 4 °C. Primary antibodies incubation followed by wash and secondary antibody incubation (1:2000 Rabbit IgG secondary, Rockland 611-1322-0500, or 1:5000 Mouse IgG secondary, Invitrogen 31430) at room temp for 1.5 h. The blots were washed with 1X TBST and developed with Western Blot Luminol Reagent (Santa Cruz Biotechnology, sc-2048). The images were quantitated using ImageJ^®^.

### Enzymatic assay

Asp-NAT activity of NAT8L was assayed radiochemically in a mixture (200 μL final volume) comprising of 10 mM potassium phosphate, 20 mM HEPES, pH 7.4, 1 mM MgCl_2_, 50 μM L-aspartate, 10000 cpm L-[U-^14^C] aspartate (Moravek Inc.), containing approximately 20 mg of protein/ mL. After incubating 30 min at 30 °C, the reaction was stopped by a 5-min incubation at 80 °C and 1 mL of 5 mM HEPES, pH 7.4, was added. The sample was centrifuged for 5 min at 13,200 rpm and the supernatant was applied on to a 1 mL Dowex AG1×8 column (Cl^−^ form, 100 – 200 mesh; Acros Organics) prepared in a Bradford column. The latter was washed with 5 mL of 5 mM HEPES, pH 7.0, followed by 5 mL of 100 mM NaCl in the same buffer, to elute unreacted aspartate followed by 5 mL of 300 mM NaCl, to elute the NAA. Radioactivity was measured using a liquid-scintillation counter.

### NAA measurement

Level of NAA were measured in neuronal cells by HPLC. A Whatman Partisil 10 SAX anion-exchange column (4.6 mm × 250 mm) was used in an Agilent 1100 Series HPLC Value System. The mobile phase consisted of 0.1 M KH_2_PO_4_ and 0.025 M KCl at pH 4.5. A flow rate of 1.5 mL/min, and the flow was monitored with an Agilent 1100 series UV detector at 214 nm. The retention time was determined based upon the retention time of an NAA standard (Sigma). Peak areas were acquired with Agilent Chemstation^®^ software.

### Design of a vector expressing *NAT8L* mRNA and mRNA oxidation

Human NAT8L expressing plasmid was purchased from applied biomaterials. Messenger RNA was obtained by *in vitro* run off transcription. The purified mRNA was oxidized using Fenton chemistry-RNA was incubated in 100 µM freshly prepared H_2_O_2_ and 10 µM FeSO_4_ at 37 ºC for an hour. The reaction was quenched using 100 µM deferoxamine mesylate (DFOM) and passed through G-25 columns (GE Healthcare) followed by denaturing polyacrylamide gel purification. The amount of purified RNA was spectrophotometrically quantified, and the total amount of RNA was normalized to that in the control reaction before being used for *in vitro* translation.

### *In vitro* translation

To translate mRNAs encoding for NAT8L protein, nuclease treated rabbit reticulocytes system (Promega) was used according to the manufacturer’s protocol. All reactions were performed in a volume of 50 μL at 30 °C for 60 min. Relative quantification of protein thus produced was performed by comparing the relative band intensities using western blot protocol as described above.

### Statistical analysis

If not otherwise stated, results are mean values ± SD of at least three independent experiments, or results show one representative experiment of a minimum of three. Statistical analyses were performed on all available data. Unless otherwise mentioned, statistical significance was determined using the two-tailed Student’s t test with p values ≤ 0.05 considered statistically significant.

## Data availability

The data that support the findings of this study are available within this manuscript and SI appendix. RNA-seq data can be accessed in Gene Expression Omnibus database (GSE144294).

## Author contributions

S.B., J.M., and P.K. conceived the project and S.B. supervised the project. P.K. designed and undertook the cellular, biochemical, and molecular biology experiments. P.K., N.W., and N.S. undertook the animal and postmortem tissue works. N.S. performed HPLC analyses and NAA measurement. A.C. and P.K. performed the RNA-seq data analysis and downstream pathway analysis. P.K., N.W., J.R. and L.S. performed the molecular biology experiments. E.F. participated in discussion with S.B, P.K., and J.M., and S.B., J.M., and, P.K. wrote the manuscript.

## Acknowledgements

We would like to thank the Rocky Mountain MS Center and the Human Brain and Spinal Fluid Resource Center at UCLA for MS and non-MS postmortem brain tissue. The research was supported from Kent State University. PK acknowledges partial research support from Graduate Student Senate, Kent State University.

