## Supplementary material for "*NAT8L* mRNA oxidation is linked to neurodegeneration in multiple sclerosis"

**This PDF file includes:**

Supplementary text

Figures S1 to S5

Tables S1 to S2

**Other supplementary materials for this manuscript include the following:**

Datasets S1 (Gene Expression Omnibus database)

**Supplementary material**

1. **Demographic information of the tissue donors**

**Table S1**

| Brain regions used in this study | Donor ID | Age | Gender | Postmortem Interval | |
|---|---|---|---|---|---|
| Cortex | 1 | 52 | F | 20.6 | MS donors |
| | 2 | 59 | M | 15.0 | |
| | 3 | 63 | F | 23.0 | |
| Cortex | 4 | 73 | F | 12.0 | Non-MS donors |
| | 5 | 75 | F | 15.4 | |
| | 6 | 59 | M | 20.2 | |

2. **List of primers used in this study**

**Table S2**

| | |
|---|---|
| Human NAT8L forward primer | 5’-TGTGCATCCGCGAGTTCCGT |
| Human NAT8L reverse primer | 5’-CGGAAGGCCGTGTTAGGGAT |
| Mouse NAT8L forward primer | 5’-TGTGCATCCGCGAGTTCCGC |
| Mouse NAT8L reverse primer | 5’-GCGGAAAGCCGTGTTGGGGA |
| Human GAPDH forward primer | 5’- GTCTCCTCTGACTTCAACAGCG |
| Human GAPDH reverse primer | 5’- ACCACCCTGTTGCTGTAGCCAA |
| Mouse GAPDH forward primer | 5’- CATCACTGCCACCCAGAAGACTG |
| Mouse GAPDH reverse primer | 5’- ATGCCAGTGAGCTTCCCGTTCAG |
| Human SOX4 forward primer | 5’-GCACATGGCTGACTACCCC |
| Human SOX4 reverse primer | 5’-GCCTTGTACAGCGAGTGGT |

| | |
|---|---|
| Human Caspase-1 forward primer | 5′-GCCTGTTCCTGTGATGTGGAG |
| Human Caspase-1 reverse primer | 5′-TGCCCACAGACATTCATACAGTTTC |
| Human COX7A1 forward primer | 5’-TGTGGCAGAGAAGCAGAAG |
| Human COX7A1 reverse primer | 5’-AGCCCAAGCAGTATAAGCAG |
| Human β-Actin forward primer | 5’-CACCATTGGCAATGAGCGGTTC |
| Human β-Actin reverse primer | 5’-AGGTCTTTGCGGATGTCCACG |

#### 3. Relative quantitation of RNA oxidation in SNP stressed SH-SY5Y cells

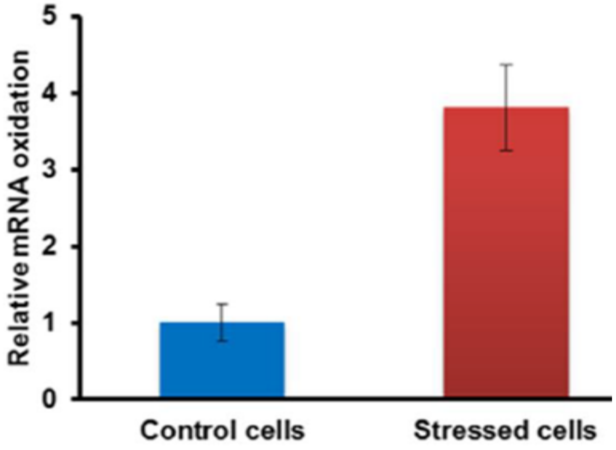


Figure S1. Relative change in mRNA oxidation in the control versus SNP stressed cells. The immunoprecipitated (with anti 8-OG antibody) RNA was spectrophotometrically quantified. Oxidized RNA from the control cells was normalized to 1.

### 4. PANTHER pathway analysis of differentially expressed transcripts

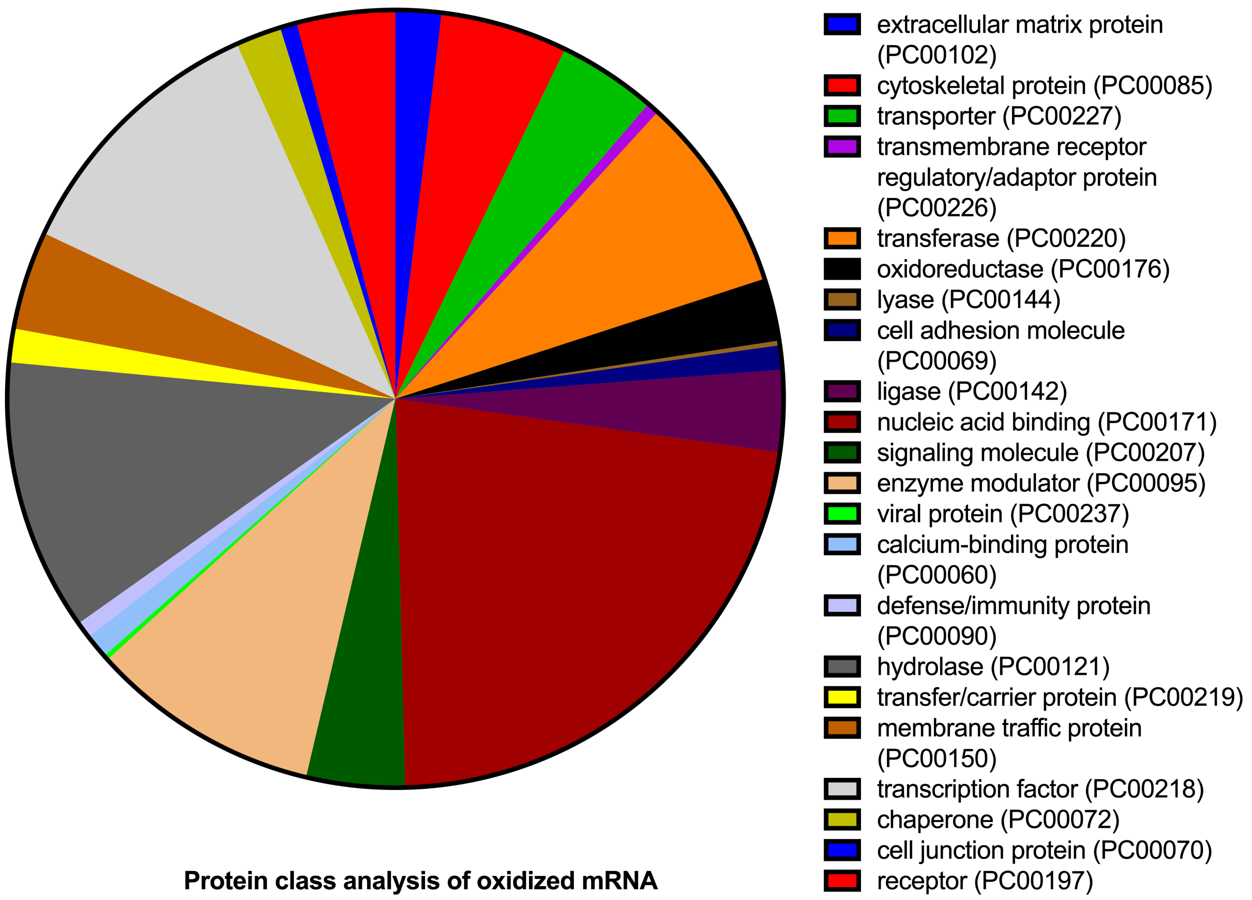


Figure S2. i. Transcript analysis based on protein class reveals that many of the mRNAs for nucleic acid binding proteins and transcription factors are selectively oxidized in human neurons under SNP induced oxidative stress.

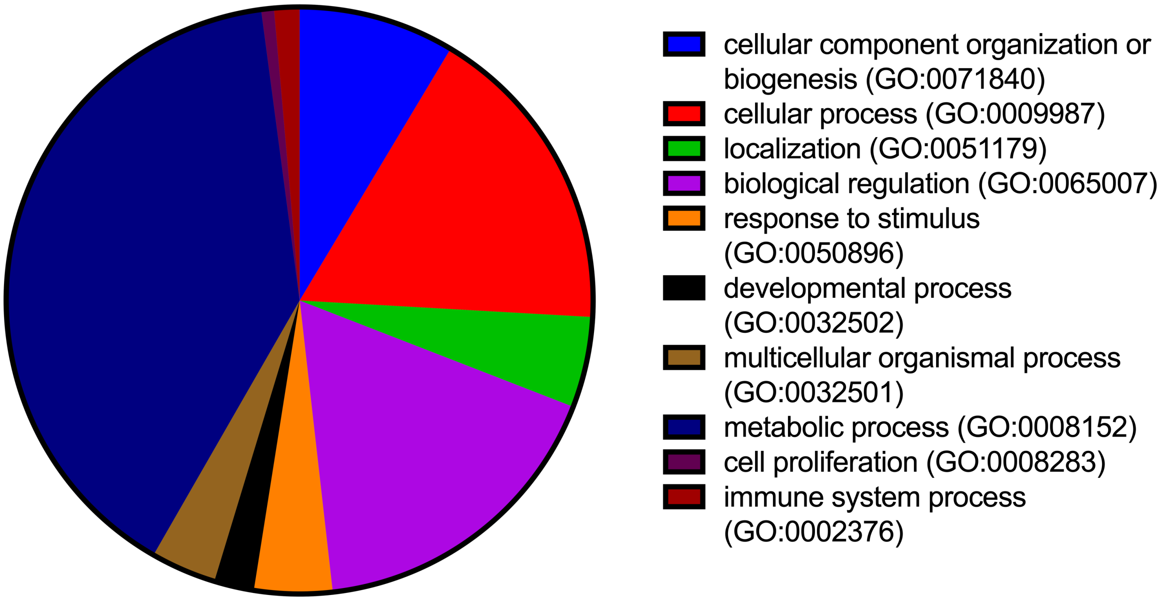


**Biological processes analysis of oxidized mRNA**

Figure S2. ii. Transcript analysis based on biological processes reveals that many of the mRNAs for metabolic pathways and cell process pathways are selectively oxidized in human neurons under SNP stress.

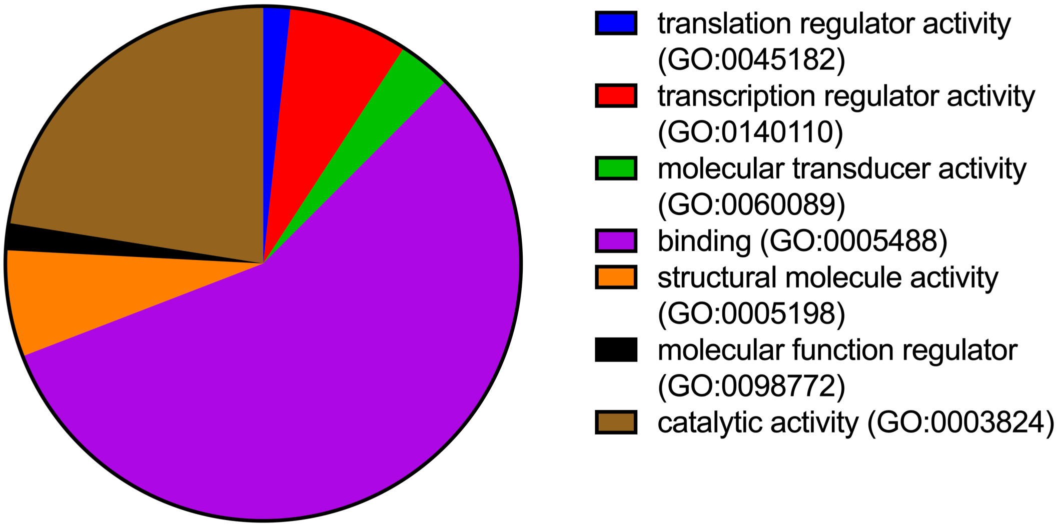


**Molecular function analysis of oxidized mRNA**

Figure S2.iii. Transcript analysis based on molecular function reveals that many of the mRNAs for binding proteins and catalytic proteins are selectively oxidized in human neurons under SNP stress.

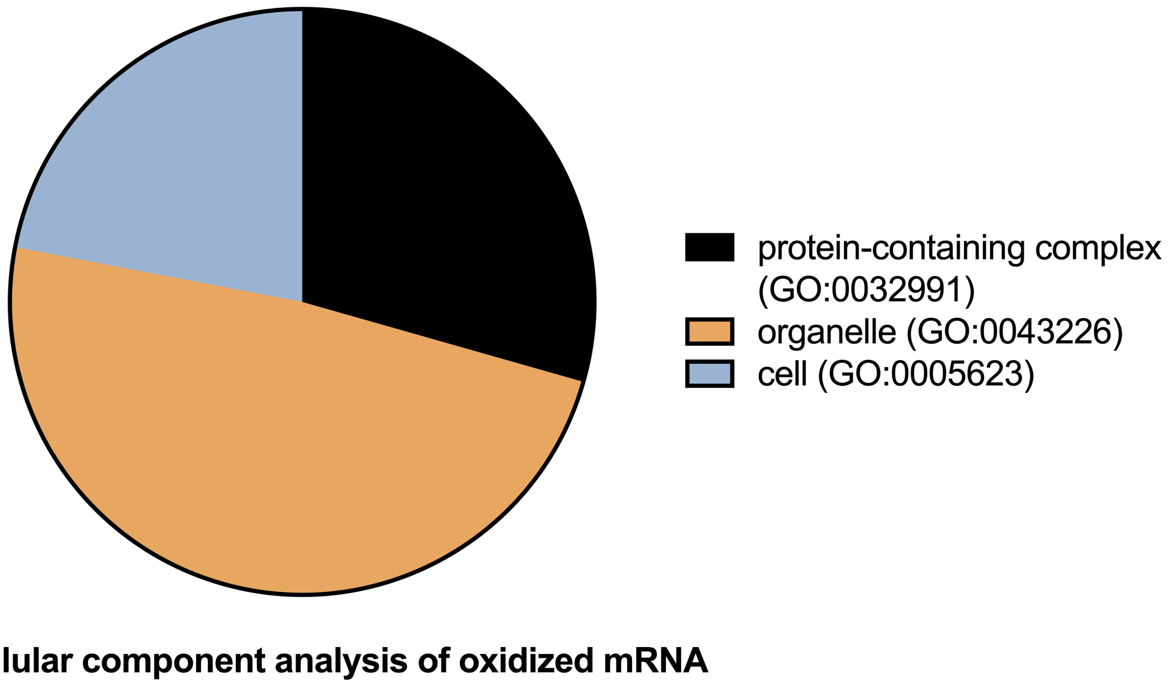


Figure S2.iv. Transcript analysis based on cellular components reveals that mRNAs encoding proteins for different cellular compartments are selectively oxidized in human neurons under SNP stress.

### 5. NAT8L protein is expressed in the neuronal cells but not in the glial cells in mouse brain

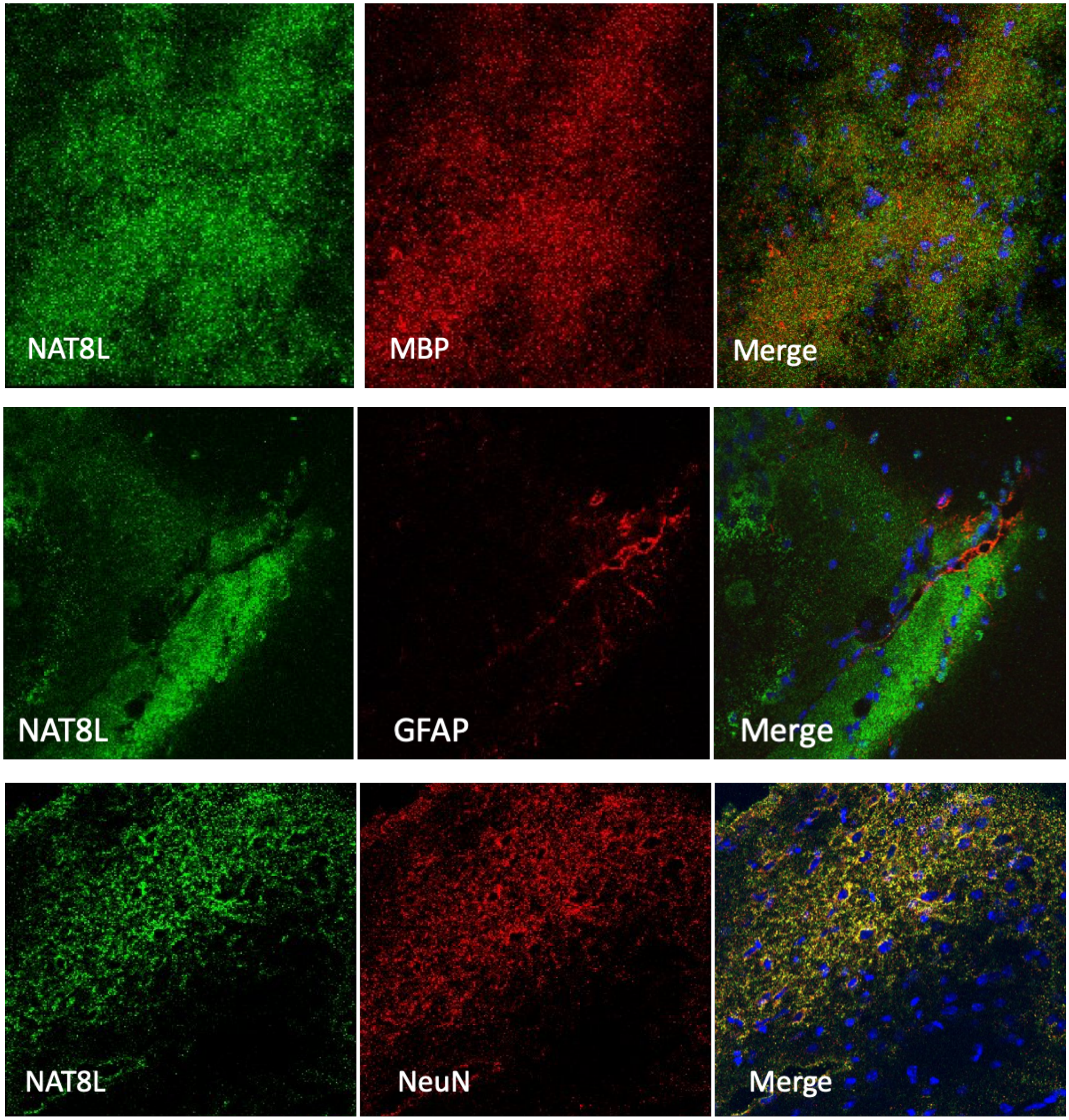


Figure S3. NAT8L protein is almost exclusively present in the neuronal cells of mouse brain. Top panel- Comparison of NAT8L stain with MBP stain (oligodendrocyte marker), middle panel- Comparison of NAT8L stain with GFAP stain (astrocyte marker), and bottom panel- Comparison of NAT8L stain with NeuN stain (neuronal marker).

6. **Asp-Nat enzymatic activity in cuprizone-fed mice brain is lowered**

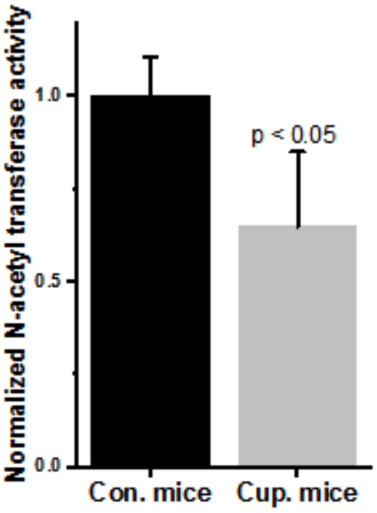


Figure S4. NAT8L enzymatic activity is compromised in cuprizone fed mice brain

7. **Selective mRNA oxidation does not seem to be a function of a higher guanosine (G) density in the transcripts (data presented in supplementary excel file 1)**

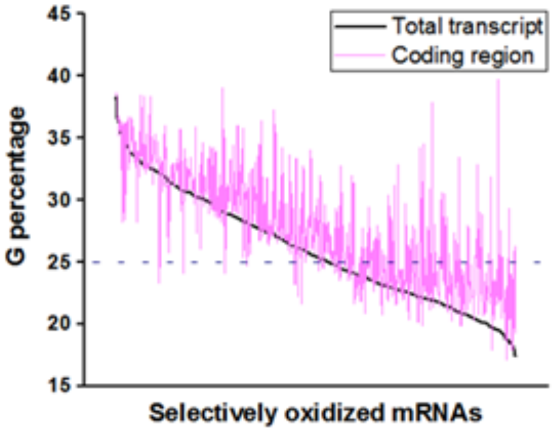


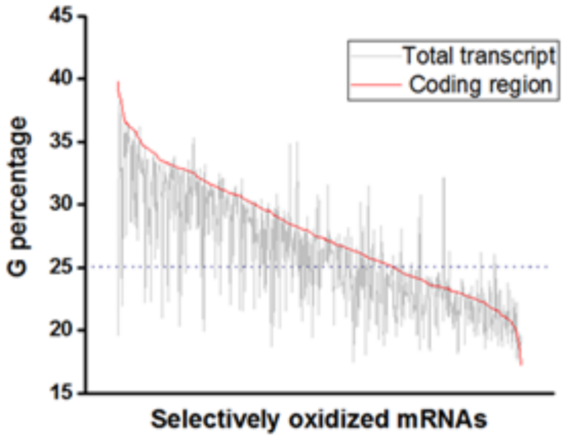


Figure S5. The selective oxidation of mRNA as a function of G enrichment in the transcripts and coding regions. Left. A scatter plot showing the trend of G% in oxidized transcripts (arranged based on decreased number of Gs in the total transcript). Right. A scatter plot showing the trend of G% in oxidized transcripts (arranged based on decreasing number of Gs in the coding region).
